# Climate variability and the paradox of plasticity

**DOI:** 10.64898/2026.08.24.746805

**Authors:** Alex R. Gunderson, Michael L. Logan, Guillermo Garcia-Costoya

**Author notes:** **Email addresses:**.

## Abstract

Adaptive phenotypic plasticity is expected to evolve when environmental conditions change predictably over time. This has led to the hypothesis that ectotherms in environments with low temperature seasonality, such as the tropics, should evolve lower thermophysiological plasticity than those from more seasonal environments (the Climate Variability Plasticity Hypothesis, or CVHP). Yet, empirical support for the CVHP is incredibly low, creating a need to identify other factors that can help explain how thermal plasticity evolves. Here, we use numerical models to show that the evolution of constitutive thermal tolerance breadth greatly affects the evolutionary benefits of thermal plasticity. In particular, tolerance breadth interacts with within- and between-season temperature variation in ways that can confound expectations of the CVHP, including conditions in which organisms from less seasonal environments benefit most from expressing plasticity. Our findings indicate that a more holistic view of the relationship between thermophysiology and environmental temperature is needed to explain the evolution of thermal plasticity across climatic gradients.

## Introduction

Phenotypic plasticity, the capacity for organisms to express different phenotypes in different environments, is a ubiquitous feature of life. Phenotypic plasticity plays a central role in individual and population responses to global change because it enables individuals to resist the drop in fitness that would otherwise occur as their local environment shifts. Understanding when and how phenotypic plasticity evolves is therefore of both basic and practical importance. In general, adaptive phenotypic plasticity is expected to evolve when environmental conditions fluctuate predictably over time (Levins, 1963; Lande, 2009; Reed et al., 2010). This has led to the hypothesis that plasticity in thermophysiological traits, such as tolerance and sensitivity to heat and cold, is greatest in regions with high levels of temporal temperature variation (hereafter referred to as the Climate Variability Plasticity Hypothesis, or CVPH). The CVPH predicts that, for example, ectotherms in the tropics, where there is little seasonal temperature change, should evolve lower thermophysiological plasticity than those at high latitudes, where temperature seasonality is greater (Janzen, 1967; Ghalambor et al., 2006; Angilletta, 2009).

Because the CVPH is intuitive, it remains a central hypothesis in ecological and evolutionary physiology. Yet, paradoxically, analyses across phylogenetic and geographic scales have repeatedly failed to uncover a consistent positive relationship between plasticity in thermal physiology and latitude (Brattstrom, 1968; Brattstrom, 1970; Angilletta, 2009; Gunderson & Stillman, 2015; Campos et al., 2021; Weaving et al., 2022). Others have found mixed support, with expected patterns found in some subsets of data (e.g., some taxa or thermal traits) but not others (Rohr et al., 2018; Morley et al., 2019), including patterns opposite to the CVPH, with taxa at higher latitudes actually exhibiting less plastic capacity (Seebacher et al., 2015; Rohr et al., 2018). Latitude alone does not perfectly capture geographic differences in temporal temperature variation, so analyses have also been conducted that directly test for links between thermal plasticity and environmental temperature variation (e.g., temperature seasonality). These analyses have similarly found no support for the CVPH (Comte & Olden, 2017), mixed support (Gunderson & Stillman, 2015; Rohr et al., 2018), and inverse patterns (Rohr et al., 2018; Ruthsatz et al., 2024). The cause of this profound lack of correspondence between theoretical prediction and empirical reality is a mystery and indicates that we must look into forces beyond temporal climate variability to fully understand the evolution of plasticity in thermal physiology.

An important, though often overlooked, consideration when examining the evolution of thermal plasticity is that plasticity is not the only means of dealing with fluctuating environmental conditions. Rather than evolving an induced plastic response, organisms can evolve a constitutively expressed phenotype that enables high performance over a broad range of conditions (Lynch & Gabriel, 1987; Scheiner, 1993). Because both plasticity and constitutive tolerance breadth can serve as solutions to the same problem, our expectations for the evolution of plasticity should depend not just on the scale and predictability of environmental variation, but on the constitutive tolerance breadth of the organisms under consideration.

In ectotherms, constitutive thermal tolerance breadth frequently evolves predictably across latitudinal and other thermal gradients, with animals that live in more variable environments evolving wider breadths (Deutsch et al., 2008; Huey et al., 2009; Sunday et al., 2011; MunEoz et al., 2014; Sheldon & Tewksbury, 2014; Buckley & Huey, 2016; Gutiérrez–Pesquera et al., 2016; Payne & Smith, 2017; Shah et al., 2017; von May et al., 2017; Dewenter et al., 2024; Dewenter et al., 2025; Yilmaz et al., 2026). We propose that this systematic variation in constitutively expressed tolerance breadth could be one key to explaining the lack of empirical support for the CVPH. In what follows, we use numerical models to demonstrate how the evolutionary benefits of plasticity in thermal physiology depend crucially on constitutive thermal tolerance breadth in ways that can confound expectations of the CVPH.

### Constitutively expressed thermal tolerance breadth and the benefits of plasticity

Consider three ectothermic organisms with stereotypically asymmetrical thermal performance curves (TPCs; Sinclair et al., 2016) that differ in breadth (Tbr) within a realistic range (e.g., Deutsch et al., 2008; Sunday et al., 2011; Sheldon & Tewksbury, 2014; von May, 2017; see other citations above). We refer to these as narrow (Tbr = 15 °C), intermediate (Tbr = 30 °C), and wide (Tbr = 45 °C) TPCs (Figure 1). Assume that each organism’s starting body temperature (Tb) is constant and at the temperature that maximizes fitness (thermal optimum, Topt). Now let’s assume the body temperature of each organism increases by an identical 3 °C, representing seasonal warming (Figure 1, gray vs. orange vertical line and arrow). In the absence of phenotypic plasticity, fitness declines for all organisms (Figure 1, gray vs. black horizontal lines). The magnitude of this fitness decrement is dependent on Tbr, with the narrowest TPC experiencing the greatest fitness decline and the widest TPC experiencing the least fitness decline (see also Deutsch et al., 2008; Huey et al., 2009).

**Figure 1:**
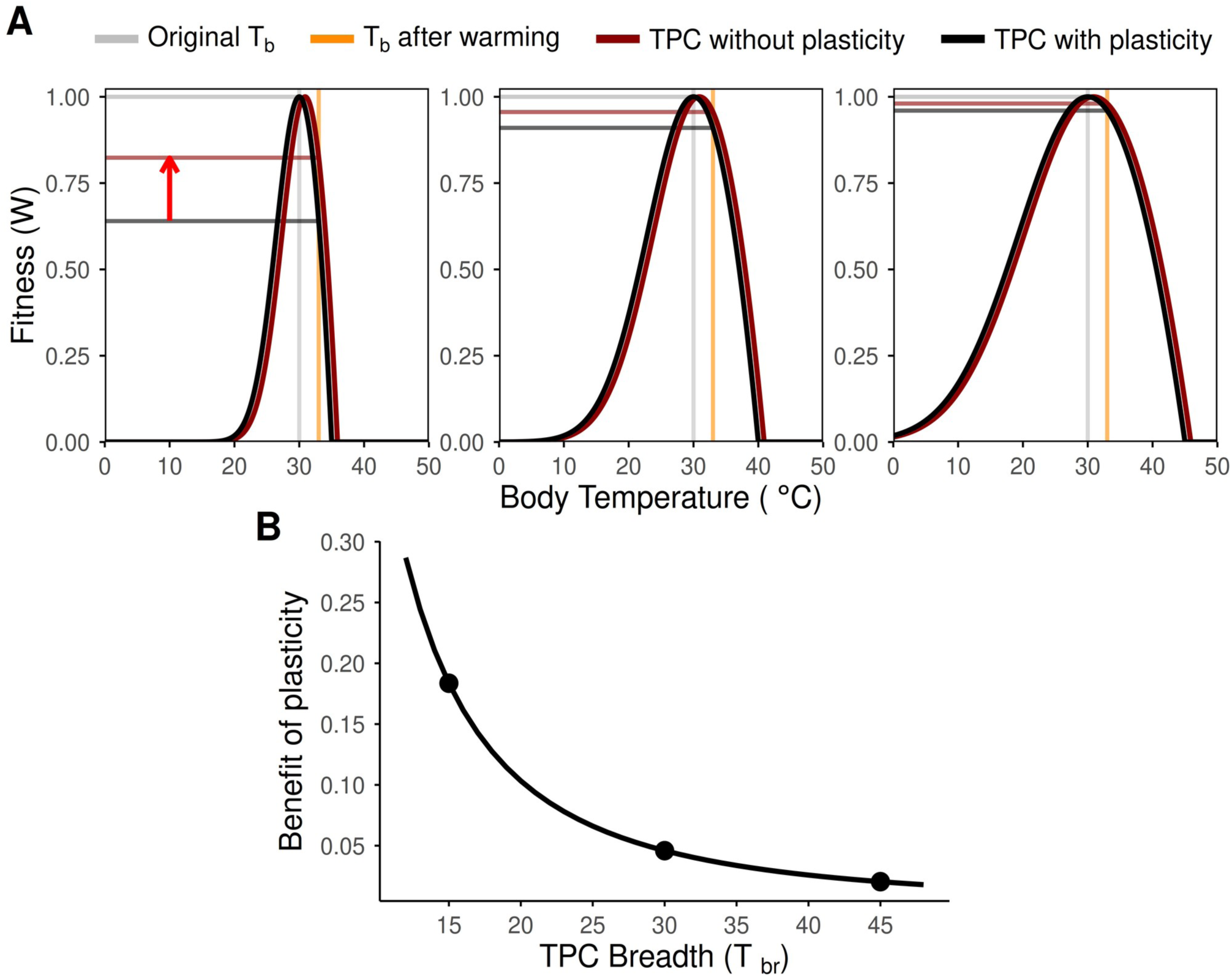
Phenotypic plasticity can provide greater net fitness benefits for narrow than for broad thermal performance curves (TPCs). Panel A shows three organisms with TPCs that vary in their constitutive thermal breadth (Tbr): narrow (Tbr = 15 °C), left panel, intermediate (Tbr = 30 °C, middle panel), and wide (Tbr = 45 °C, right panel). The organisms have evolved to maximize fitness (W = 1) at the constant body temperature they experience, which in this case we set arbitrarily to 30 °C (Topt, gray vertical line). Following a 3 °C increase in body temperature (orange vertical line), absolute fitness declines across all organisms in the absence of plasticity (projected onto the y-axis via the gray vs. black horizontal lines), with the narrowest TPC experiencing the most severe decline in fitness. If all three organisms shift their TPC in response to warming through phenotypic plasticity by the same amount (here assuming an Acclimation Response Ratio or ARR of 0.30), the net fitness benefit conferred by plasticity (the vertical distance between the red and black horizontal lines, indicated by the red arrow on the top left panel) is highest for the organism with the narrowest TPC and lowest in the organism with the widest TPC. Panel B shows how the benefit of plasticity declines exponentially with increasing TPC breadth, with the three dots indicating the three TPC breadths shown in panel A.

Next, we examined what happens when these organisms undergo a plastic shift in response to the higher body temperature (Figure 1, red curves). Empirically, thermal plasticity does not yield perfect compensation (Seebacher et al., 2015; Gunderson & Stillman 2015). Acclimation Response Ratios (ARRs), which describe how features of TPCs shift for each °C change in Tb, generally range between 0.1 and 0.5 (Gunderson & Stillman, 2015), meaning that TPC features shifts 0.1–0.5 °C higher for every 1 °C increase in Tb. We assume that each organism exhibits an identical ARR of 0.3 for this and all subsequent analyses.

After plastic adjustment, fitness under the warmer regime is improved for all taxa relative to the state without plasticity (Figure 1, red vs. black horizontal lines), but still declines commensurate with TPC breadth because plasticity is not fully compensatory (Figure 1, red vs. gray horizontal lines). Two other key patterns emerge. First, the net fitness benefit of plasticity (i.e., the difference in fitness with and without plasticity; red versus black horizontal lines in Figure 1) is highest for the organism with the narrowest TPC and lowest for the organism with a wider TPC. In other words, narrow TPCs do not just make fitness more sensitive to changes in body temperature; they can also make fitness more sensitive to plastic shifts in the TPC itself.

Second, despite achieving the highest net benefit of plasticity, the narrow TPC organism’s absolute fitness post-plastic shift remains lower than that of its intermediate and wide TPC counterparts (Figure 1). Therefore, in the absence of complete plastic compensation, organisms with narrow TPCs must have greater plastic capacity than organisms with broad TPCs to maintain equivalent absolute fitness under the same level of environmental change. In our example, the narrow and intermediate TPC organisms would need to express ARRs of 0.76 and 0.54 to match the fitness of the wide TPC organism expressing an ARR of 0.30 (Figure 2). Note also that the benefit of evolving greater plasticity (i.e., increasing ARR) is highest for the narrow TPC organism and lowest for the wide TPC organism (Figure 2). This can be seen as a difference in the slope of the relationship between plasticity benefit and ARR in Figure 2. Therefore, under similar environmental conditions, the benefit of evolving greater plasticity increases when constitutively expressed TPCs are narrower.

**Figure 2:**
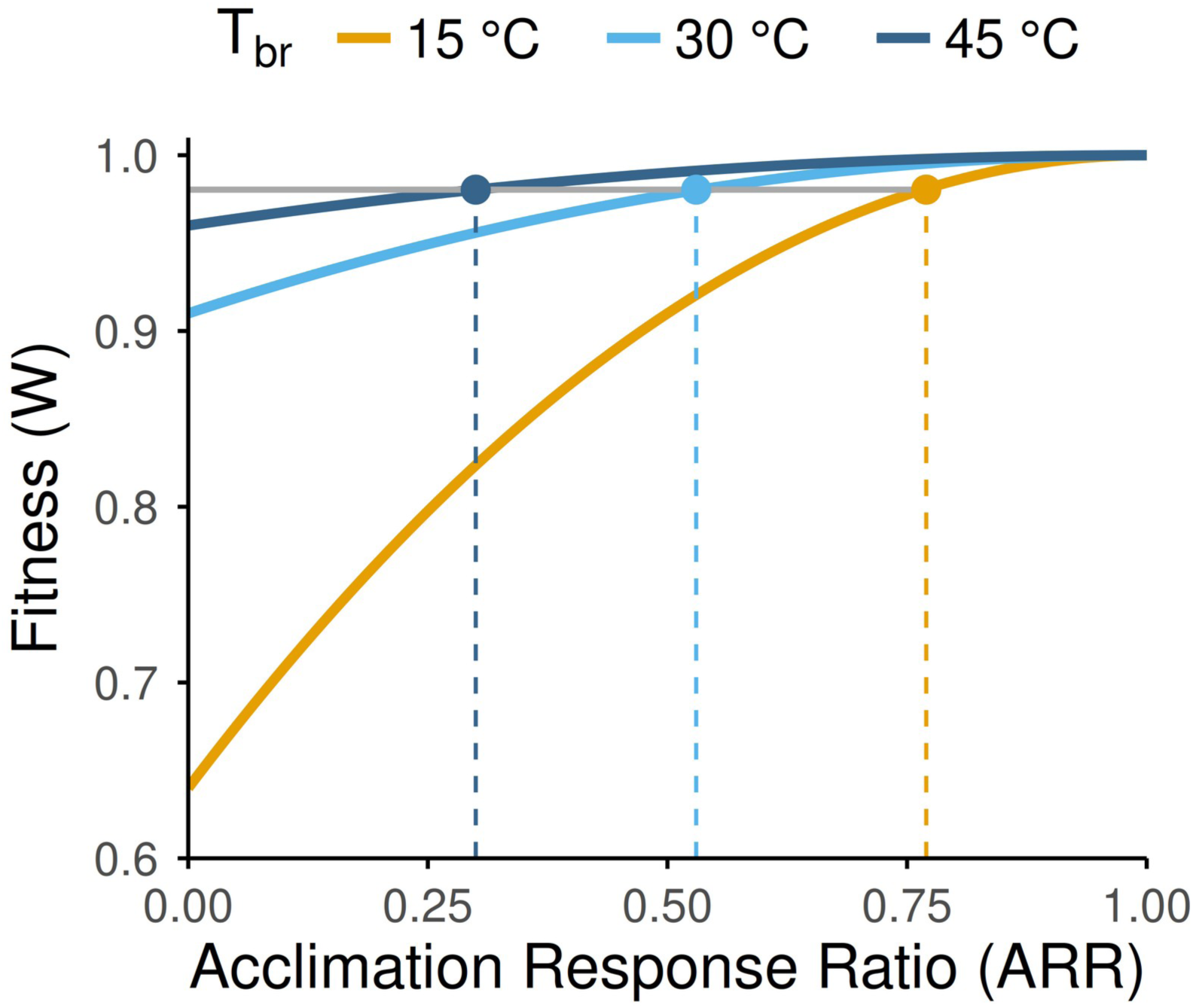
Organisms with narrow thermal performance curves (TPCs) must exhibit greater plasticity to match the fitness of organisms with broad TPCs after warming. Results here are from the scenario described in Figure 1, in which taxa with different TPC breadths experience a 3 °C increase in body temperature (Tb). The organisms with the narrow (15 °C) and intermediate (30 °C) TPC would need to exhibit Acclimation Response Ratios (ARRs) of 0.77 and 0.54, respectively (yellow and light blue lines) to maintain the same level of fitness as the organism with the wide TPC (45 °C) exhibiting an ARR of 0.30 (dark blue lines).

### Incorporating within-season variability in body temperature

We now build on the model by adding variation around the mean body temperature experienced at each time point, represented by the standard deviation of body temperature, 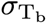 (Figure 4). This can be thought of as, for example, variation in body temperature experienced within a season, and thus we refer to it as “within-season body temperature variation.” As before, we assume that these organisms have evolved to maximize initial fitness, which results in each taxon having a different initial mean body temperature (Figure 3A). This is because TPCs are asymmetrical, which causes fitness to be maximized when mean Tb < Topt in the presence of thermal variability (i.e., Jensen’s inequality; Denny, 2017; Martin & Huey, 2008; Ruel & Ayres, 1999). Within-season body temperature variation also reduces the maximum expected fitness that organisms can achieve (see also Vasseur et al., 2014), and the effect is greatest for organisms with narrow TPCs (Figure 3).

**Figure 3:**
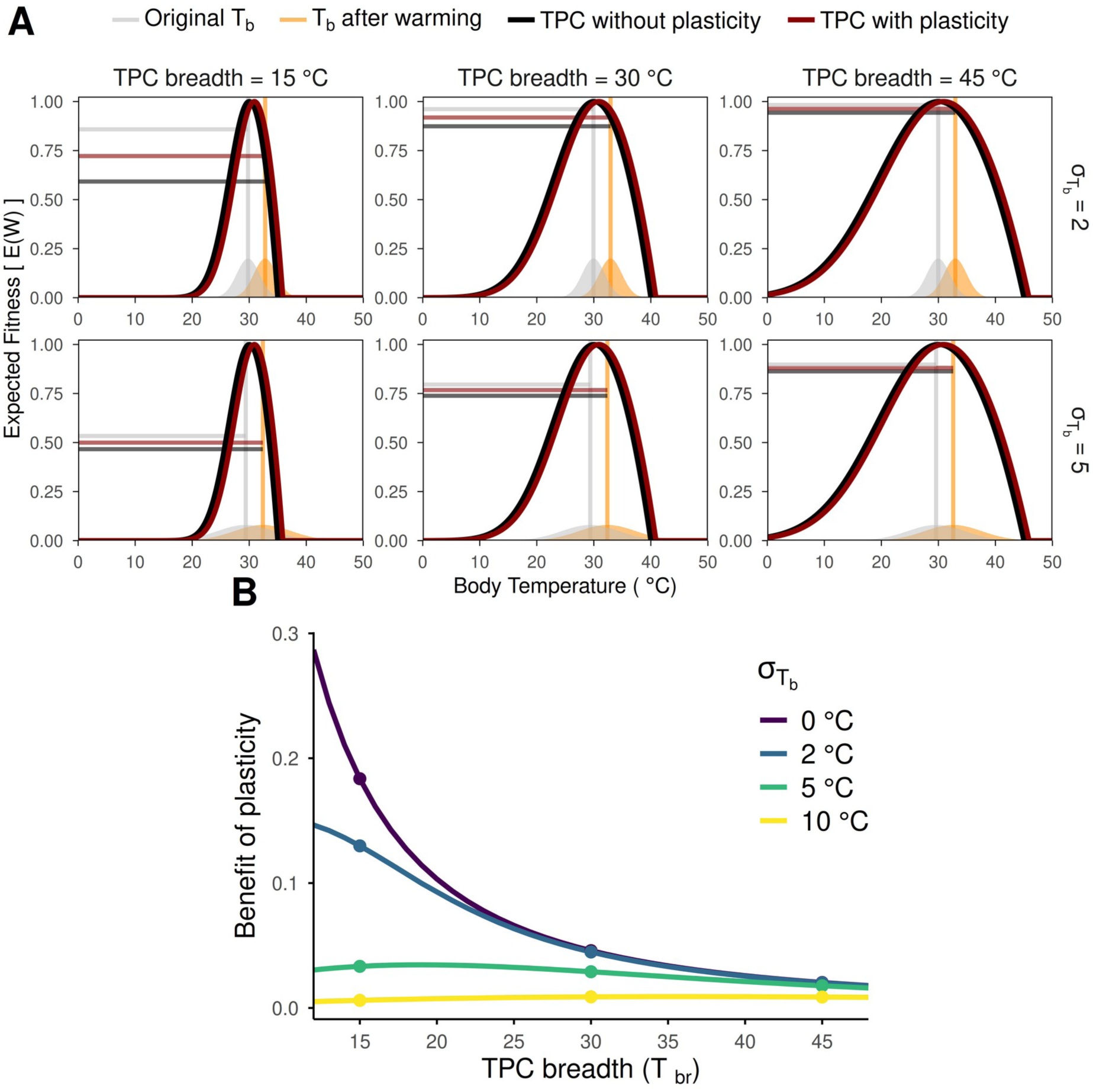
Examples of the effect of constitutive thermal breadth and within-season body temperature variation on the benefits of plasticity. (A) Thermal performance curve (TPC) breadth and within-season variation in the distribution of body temperatures, represented by the standard deviation (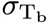), interact to determine plasticity benefits. Note that under variable within-season body temperatures, initial fitness (gray horizontal lines) is not 1 due to Jensen’s inequality (see text). After a 3 °C increase in mean body temperature (gray vs orange vertical lines and areas), the organisms shift their TPCs with an acclimation response ratio (ARR) of 0.30. The expected fitness benefit of plasticity (difference between the red and black horizontal lines) is greatest for the narrowest TPC, but for all TPCs, it decreases when body temperatures are more variable. (B) The expected benefit of plasticity as a function of both TPC breadth and within-season body temperature variation, with 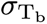 set to 0 as in Figure 1, as well as 2, 5, or 10 °C. Dots in this panel indicate TPCs with Tbr = 15, 30, and 45 °C, as shown in panel A.

**Figure 4:**
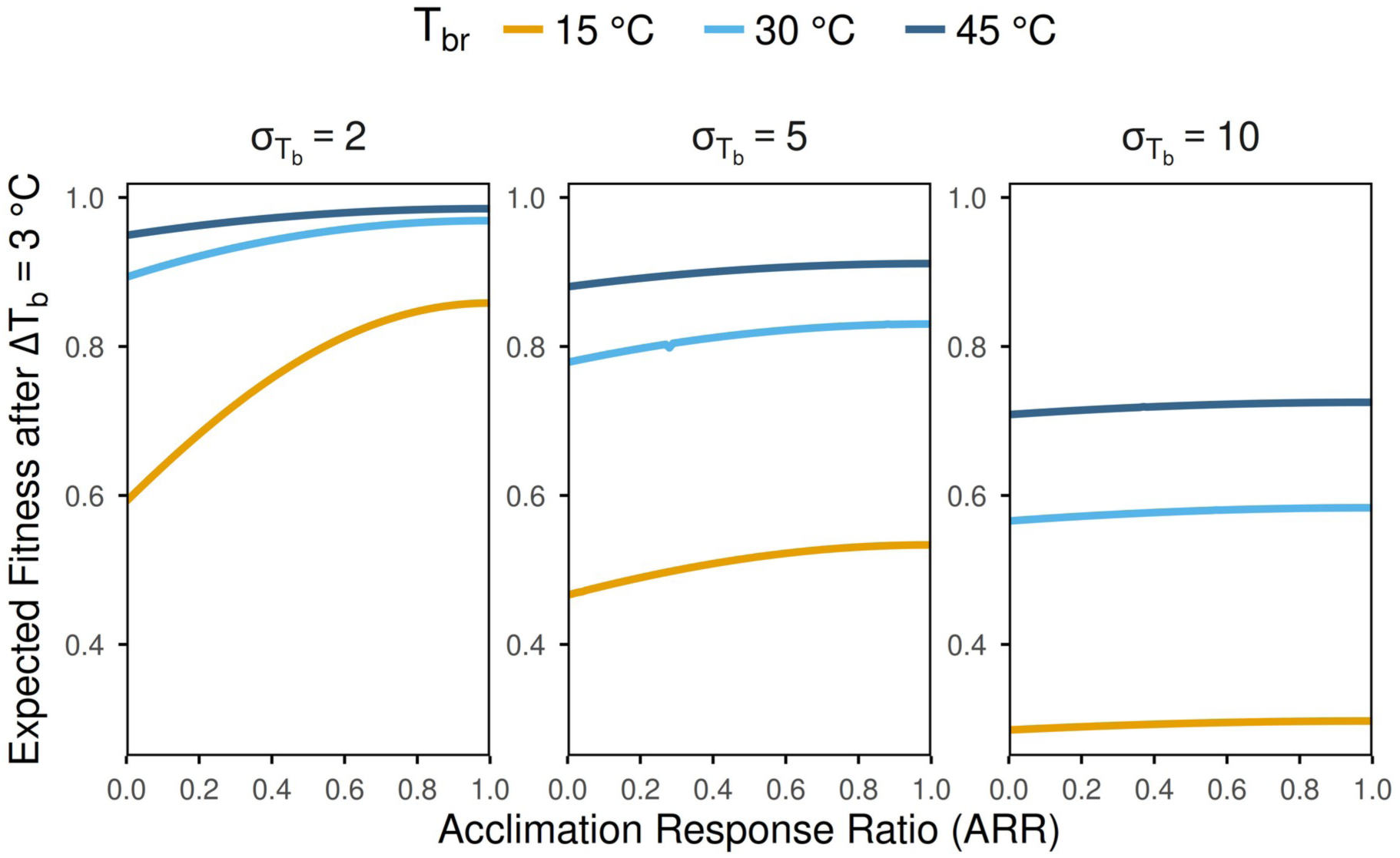
Effects of constitutively expressed thermal performance (TPC) breadth (Tbr) and within-season temperature variation (body temperature standard deviation, 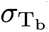 on the benefits of thermal plasticity across a range of plasticity levels (Acclimation Response Ratios, ARR). Results are based on a mean body temperature increase of 3 °C. Relatively stable within-season body temperatures (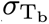 = 2 °C, left panel) lead to greater benefits of plasticity than more variable within-season body temperatures (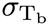 = 5 or 10 °C, center and right panel, respectively) regardless of TPC breadth. In addition, when within-season body temperatures are more stable, the benefit of evolving greater plasticity (i.e., higher ARR) is larger, and especially for narrow TPCs (compare steepness of the colored lines between values and Tbr).

If we again assume an increase in mean body temperature of 3°C, several patterns of note emerge. First, the benefit of plasticity decreases as within-season body temperature variation increases for all organisms (Figure 3). This is consistent with previous theoretical treatments with symmetrical tolerance curves (Gabriel & Lynch, 1992), and means that, for two organisms with the same thermal tolerance breadth, the one with a more stable within-season body temperature distribution will benefit most from thermal plasticity.

Second, the extent to which within-season body temperature variation reduces the benefits of plasticity is dependent on TPC breadth. As within-season body temperature variation increases, the narrow TPC organism experiences the largest decrease in the benefit of plasticity, while the wide TPC organism is affected the least (compare points within a given TPC breadth in Figure 3B). In addition, within-season body temperature variation changes who benefits the most from plasticity. When 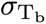 = 0, 2, or 5°C (the purple, blue, and green lines in Figure 3B, respectively), the narrow TPC organism gains more from plasticity than the wide TPC organism. However, when 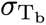 = 10°C, the narrow TPC organism benefits least, and the wide TPC organism benefits most (yellow line in Figure 3B).

### The effects of changes in mean body temperature

To this point, we have explored how TPC breadth and within-season body temperature variation influence the benefit of plasticity. We now add variation in the degree of temporal (e.g., seasonal) mean body temperature change. In general, the benefit of plasticity increases with seasonal body temperature change (Figure 5). This is consistent with the expectation that plasticity should be favored when there is greater temporal change in environmental temperature. However, when within-season body temperature variation is low (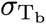 = 2 °C), the narrow TPC organism eventually experiences decreasing plasticity benefits with increasing seasonal temperature change (orange line in Figure 5, left). This occurs because temperature change becomes too large relative to TPC breadth, even with plastic compensation.

**Figure 5:**
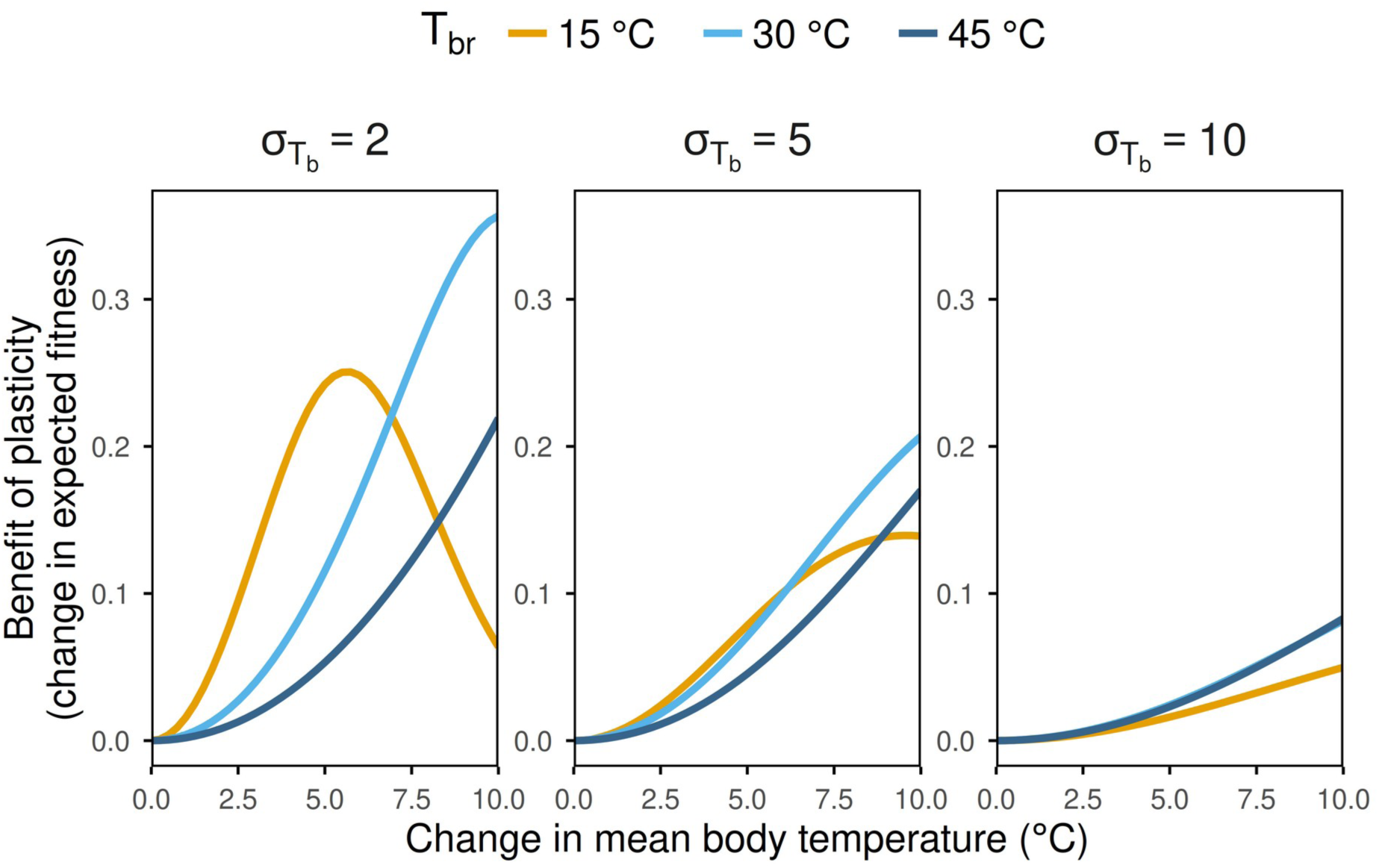
The net fitness benefit of plasticity depends on a three-way interaction between constitutively expressed thermal performance curve (TPC) breadth (Tbr), the magnitude of temporal (i.e., seasonal) body temperature change, and within-season body temperature variation (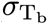).

Additionally, the relative benefit of plasticity depends on a three-way interaction between TPC breadth, within-season body temperature variation, and the magnitude of seasonal body temperature change. When within-season body temperature variation is relatively low (Figure 5, left) and the magnitude of seasonal body temperature change is relatively small (< 6 °C), the narrow TPC organism gains more fitness benefit from plasticity than the intermediate and wide-breadth organisms. However, as within-season body temperature variation increases (Figure 5 center, right), this changes. Indeed, at our highest level of within-season body temperature variation, the narrow TPC organism had the lowest benefit of plasticity across all magnitudes of seasonal temperature change (Figure 5, right). Finally, regardless of within-season body temperature variation, the narrow TPC organism received less benefit of plasticity than the intermediate and wide TPC organisms at the highest magnitudes of seasonal body temperature change (> ∼ 8 °C; Figure 5).

### Putting it all together

There are three general takeaways from our analyses. First, the benefits of thermal plasticity for ectotherms depend on constitutively expressed thermal tolerance breadth. There are conditions in which narrow TPC organisms benefit most from plasticity and conditions in which broad TPC organisms benefit most (Figures 1-5). Second, within-season body temperature variation plays a crucial role in determining the benefit of plasticity. Experiencing greater within-season body temperature variation always decreases the benefits of plasticity, but the magnitude of this effect depends on constitutive TPC breadth: organisms with narrow TPCs experience greater decreases in plasticity benefits as within-season body temperature variation increases (Figures 3, 4). Third, the magnitude of seasonal change in body temperature matters, but in a complex way that is dependent on both constitutive TPC breadth and within-season variation in body temperature. In most cases, the benefits of plasticity rise as mean seasonal body temperature change increases (Figure 5). However, the combination of low within-season body temperature variation and small changes in mean body temperature across seasons causes narrow TPC organisms to receive comparatively greater benefits of plasticity (e.g., Figure 5, left). In contrast, high within-season body temperature variation and large seasonal shifts in mean body temperature tend to yield comparatively greater plasticity benefits for wide TPC organisms (Figure 5).

Our results do not refute that the temporal temperature variation on which the CVPH rests is an important factor for the evolution of thermal plasticity (Figure 5). However, we show that temporal temperature variation cannot be considered in isolation. Constitutive TPC breadth is important, and we know that empirical TPC breadth varies predictably across climatic gradients. Ectotherms inhabiting habitats with greater temperature seasonality tend to have wider thermal tolerance breadths (e.g., Sunday et al., 2010; Sheldon & Tewksbury, 2014; Gutiérrez-Pesquera et al., 2016; Baudier et al., 2019), which may depress the benefits of plasticity by facilitating high performance across a wide range of temperatures. Within-season body temperature variation is also crucial, but is not as well understood as TPC breadth due to the difficulty of measuring field body temperatures for many organisms. That said, variation in within-season environmental temperature does change across geographic gradients. For example, within a season, high-latitude terrestrial organisms tend to experience greater variation in daily maximum air temperature than low-latitude organisms (Gunderson et al., 2017; Kingsolver & Buckley, 2017). If this pattern translates to body temperatures, within-season temperature variation would reduce the benefits of TPC plasticity more for high-latitude organisms.

Our models indicate that key features of real organisms (constitutive tolerance breadth and body temperature variation) could contribute to a lack of support for the CVPH, and even produce patterns opposite to those predicted by the CVPH (Seebacher et al., 2015; Ruthsatz et al., 2024). Most tropical organisms live in environments with little temperature seasonality, but they are also likely to have narrow TPCs and experience relatively invariant body temperature distributions, features that combine to favor the evolution of thermal plasticity. Conversely, high-latitude species often live in environments with extensive seasonal temperature change, but they also tend to have broad TPCs and relatively variable body temperatures, characteristics that would limit the benefits of plasticity.

## Conclusions & Future directions

Our results highlight areas that require greater attention to resolve outstanding questions about the evolution of thermal plasticity. First, we show that integrating information about constitutive TPC breadth would be beneficial. This likely extends to other features of TPCs that affect sensitivity, such as skewness (Buckley et al., 2022; Martin & Huey, 2008). Second, within-season body temperature variation requires much greater empirical attention with respect to the evolution of thermal plasticity. Furthermore, there may be important interactions between processes that change body temperature distributions, such as behavioral thermoregulation (Hertz et al. 1993, Logan et al., 2019; MunEoz, 2022), and the evolution of plasticity. Ultimately, our results endorse taking a more holistic view of temperature-dependent physiology and environmental temperature dynamics to deepen our understanding of how thermal plasticity evolves.

## Additional methods

In the following sections, we provide details on our numerical approach. Further, all analyses were conducted in R (v.4.6.1 ; R Core Team, 2026), using the “tidyverse” package family (Wickham et. al., 2019), as well as the “ggpubr” (v.0.6.3; Kassambara, 2026) and “viridis” (v.0.6.5. ; “Garnier et al. 2024) packages for data manipulation and visualization. All code is available at: https://ggcostoya.github.io/plasticity_benefits.

### Thermal Performance Curves (TPCs)

Across all analysis, we define the relationship between body temperature (T) and fitness (W) with the equation for a thermal performance curve (TPC) developed by Deutsch et al. (2008):

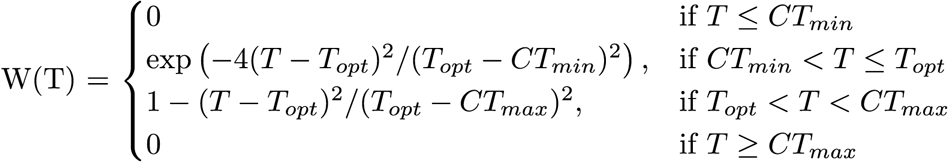

Where Topt is the optimal temperature, CTmin is the critical thermal minimum, and CTmax is the critical thermal maximum. We chose this equation because its parameters are thermal physiological traits commonly measured empirically and thus easily interpretable, and because there is no clear TPC equation that is better than any other (Kontopoulos et al. 2024). As Deutsch et al. (2008) indicate, due to the exponential nature of the ascpenting part of this TPC equation (when T < Topt), CTmin does not coresspond exactly to W = 0, but to W ∼ 0.001, although this should have minimal impact on our results. In all cases, we also assumed that Topt would be arbitrarily set at 30°C and that the asymmetry of the TPC would be set to 2/3 such that Topt - CTmin = 2 * (CTmax – Topt).

### Expected fitness calculation

To calculate the expected fitness (E[W]) of a TPC under a given body temperature distribution, we used the method developed by Vasseur et al. (2014), which involves integrating the product of the TPC and the probability density function of the body temperature distribution across all possible body temperatures using the expression:

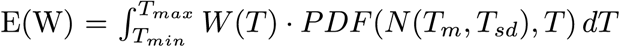

Where W(T) is the TPC function, PDF(N(Tm, Tsd), T) is the probability density function of a normal distribution with mean Tm and standard deviation Tsd, and Tmin and Tmax are the minimum and maximum body temperatures considered in the integration, which we set to - 30°C and 60°C, a sufficiently large range to encompass all possible body temperatures. For our purposes, we assume that the organism always experiences a normal distribution of body temperatures, which is a common assumption in the literature (e.g., Vasseur et al. 2014, Pinsky et al. 2019), although we acknowledge that this may not always be the case in nature.

### Optimal thermal safety margin (OTSM) calculation

To calculate the optimal thermal safety margin (OTSM) for a given TPC and body temperature distribution, we used a numerical optimization approach. We defined the OTSM as the difference between Topt and the mean body temperature (Tm) that maximizes expected fitness given the variability in body temperature (Tsd). Thus, to calculate OTSM, we needed to find the value of Tm that maximizes expected fitness for a given TPC and Tsd. To accomplish this, we defined a function that calculates expected fitness for a given Tm and Tsd using the expected fitness function described above. We then used the “optimize” function from the base R package “stats” (R Core Team, 2026) to find the value of Tm that maximizes expected fitness, which corresponds to the optimal thermal safety margin (the difference between Topt and Tm). In Supplementary Figure 1 we show how OTSM changes as a function of Tsd for TPCs of varying breadths.

## Acknowledgments

We thank James Stroud for helpful discussions and feedback on earlier versions of this manuscript and to Old Spice’s 2006 NASCAR campaign for inspiration.

## Supplementary Figures

**Supplementary Figure 1:**
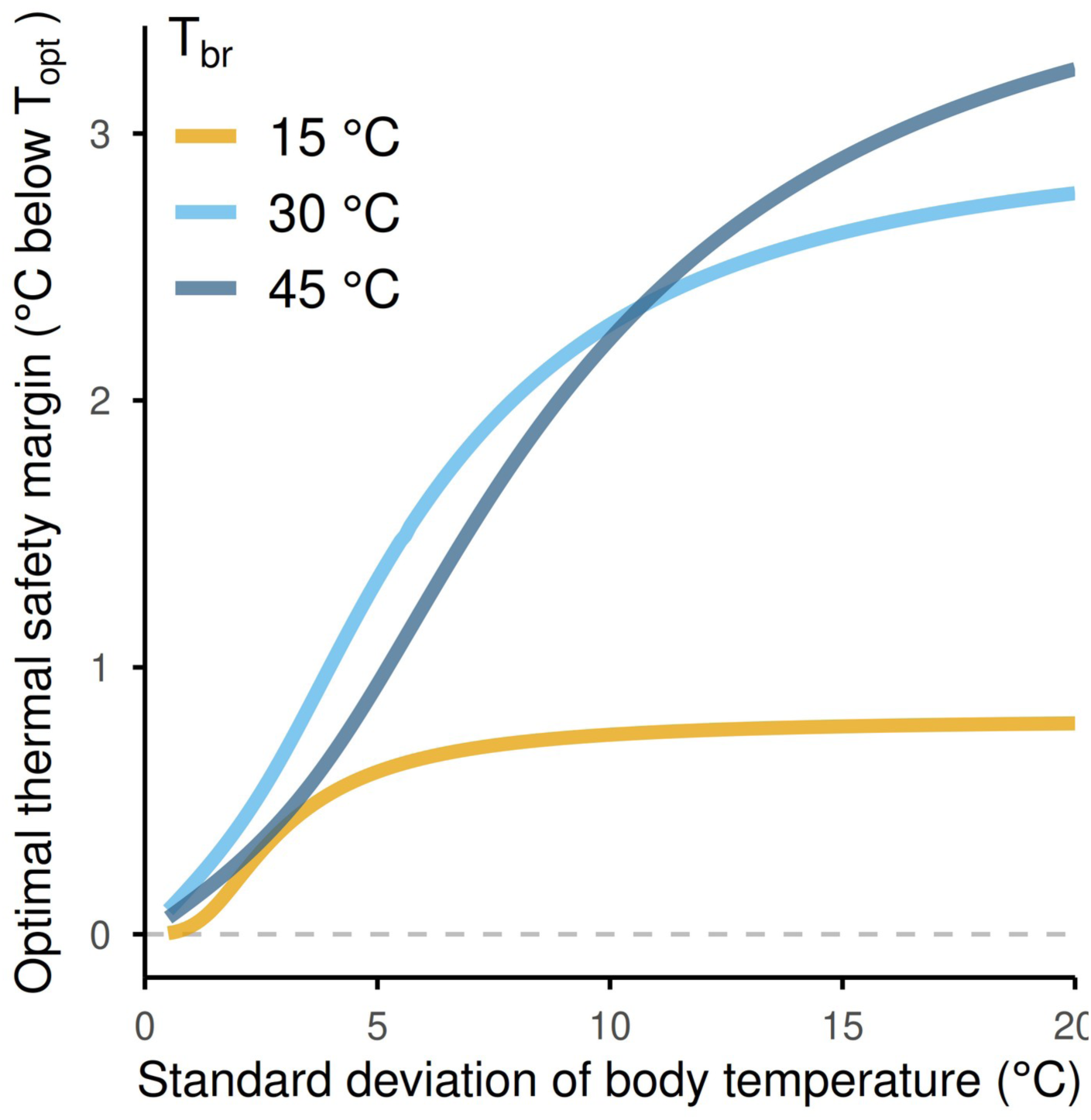
Optimal thermal safety margin (OTSM, expressed as C below a TPC’s Topt) of TPCs with varying constitutive thermal performance breadths (colored lines) as a function of the standard deviation of the body temperature distribution they experience. Note that, in all cases, OTSM increases with increasing variability in body temperature, but only up to a point where it plateaus. The value at which this stabilization occurs is greater for wider than narrower TPCs (B), and in all cases this stabilization occurs at body temperatures that are well above a TPC’s CTmin.

## References

Buckley, L. B., and R. B. Huey. 2016. Temperature extremes: geographic patterns, recent changes, and implications for organismal vulnerabilities. Global change biology 22:3829–3842.

Buckley, L. B., R. B. Huey, and J. G. Kingsolver. 2022. Asymmetry of thermal sensitivity and the thermal risk of climate change. Global Ecology and Biogeography 31:2231–2244.

Campos, D. F., R. D. Amanajás, V. M. Almeida-Val, and A. L. Val. 2021. Climate vulnerability of South American freshwater fish: Thermal tolerance and acclimation. Journal of Experimental Zoology Part A: Ecological and Integrative Physiology 335:723–734.

Comte, L., and J. D. Olden. 2017. Evolutionary and environmental determinants of freshwater fish thermal tolerance and plasticity. Global Change Biology 23:728–736.

Denny, M. 2017. The fallacy of the average: on the ubiquity, utility and continuing novelty of Jensen’s inequality. Journal of Experimental Biology 220:139–146.

Deutsch, C. A., J. J. Tewksbury, R. B. Huey, K. S. Sheldon, C. K. Ghalambor, D. C. Haak, and P. R. Martin. 2008. Impacts of climate warming on terrestrial ectotherms across latitude. Proceedings of the National Academy of Sciences 105:6668–6672.

Dewenter, B. S., D. P. Giling, A. A. Shah, J. Hughes, N. L. Poff, C. K. Ghalambor, W. C. Funk et al. 2025. Annual Temperature Range Drives Thermal Breadth of Freshwater Insects Across Multiple Spatial Scales. Ecology letters 28:e70288.

Dewenter, B. S., A. A. Shah, J. Hughes, N. L. Poff, R. Thompson, and B. J. Kefford. 2024. The thermal breadth of temperate and tropical freshwater insects supports the climate variability hypothesis. Ecology and Evolution 14:e10937.

Gabriel, W., and M. Lynch. 1992. The selective advantage of reaction norms for environmental tolerance. Journal of Evolutionary Biology 5:41–59.

Garnier, S., Noam Ross, N., Rudis, R., Camargo, A. P., Marco Sciaini, M., and Scherer, C. 2024. viridis(Lite) - Colorblind-Friendly Color Maps for R. viridis package version 0.6.5.

Ghalambor, C. K., R. B. Huey, P. R. Martin, J. J. Tewksbury, and G. Wang. 2006. Are mountain passes higher in the tropics? Janzen’s hypothesis revisited. Integrative and Comparative Biology 46:5–17.

Gunderson, A. R., M. E. Dillon, and J. H. Stillman. 2017. Estimating the benefits of plasticity in ectotherm heat tolerance under natural thermal variability. Functional Ecology 31:1529–1539.

Gunderson, A. R., and J. H. Stillman. 2015. Plasticity in thermal tolerance has limited potential to buffer ectotherms from global warming. Proceedings of the Royal Society of London B: Biological Sciences 282:20150401.

Gutiérrez-Pesquera, L. M., M. Tejedo, M. Á. Olalla-Tárraga, H. Duarte, A. Nicieza, and M. Solé 2016. Testing the climate variability hypothesis in thermal tolerance limits of tropical and temperate tadpoles. Journal of Biogeography 43:1166–1178.

Hertz, P. E., R. B. Huey, and R. Stevenson. 1993. Evaluating temperature regulation by field-active ectotherms: the fallacy of the inappropriate question. The American Naturalist 142:796–818.

Huey, R. B., C. A. Deutsch, J. J. Tewksbury, L. J. Vitt, P. E. Hertz, H. J. Á. Pérez, and T. Garland. 2009. Why tropical forest lizards are vulnerable to climate warming. Proceedings of the Royal Society B: Biological Sciences 276:1939–1948.

Janzen, D. H. 1967. Why mountain passes are higher in the tropics. The American Naturalist 101:233–249.

Kassambara, A. 2026. ggpubr:’ggplot2’based publication ready plots. R package version 0.6.3.

Kingsolver, J. G., and L. B. Buckley. 2017. Quantifying thermal extremes and biological variation to predict evolutionary responses to changing climate. Philosophical Transactions of the Royal Society B: Biological Sciences 372:20160147.

Kontopoulos, D. G., Sentis, A., Daufresne, M., Glazman, N., Dell, A. I., & Pawar, S. 2024. No universal mathematical model for thermal performance curves across traits and taxonomic groups. Nature Communications, 15(1), 8855.

Lande, R. 2009. Adaptation to an extraordinary environment by evolution of phenotypic plasticity and genetic assimilation. Journal of Evolutionary Biology 22:1435–1446.

Levins, R. 1963. Theory of fitness in a heterogeneous environment. II. Developmental flexibility and niche selection. The American Naturalist 97:75–90.

Logan, M. L., J. van Berkel, and S. Clusella-Trullas. 2019. The Bogert Effect and environmental heterogeneity. Oecologia 191:817–827.

Lynch, M., and W. Gabriel. 1987. Environmental tolerance. The American Naturalist 129:283–303.

Martin, T. L., and R. B. Huey. 2008. Why “suboptimal” is optimal: Jensen’s inequality and ectotherm thermal preferences. The American Naturalist 171:E102–E118.

Morley, S., L. Peck, J. Sunday, S. Heiser, and A. Bates. 2019. Physiological acclimation and persistence of ectothermic species under extreme heat events. Global Ecology and Biogeography 28:1018–1037.

MunEoz, M. M. 2022. The Bogert effect, a factor in evolution. Evolution 76:49–66.

MunEoz, M. M., M. A. Stimola, A. C. Algar, A. Conover, A. J. Rodriguez, M. A. Landestoy, G. S. Bakken et al. 2014. Evolutionary stasis and lability in thermal physiology in a group of tropical lizards. Proceedings of the Royal Society B: Biological Sciences 281:20132433.

Payne, N. L., and J. A. Smith. 2017. An alternative explanation for global trends in thermal tolerance. Ecology Letters 20:70–77.

Pinsky, M. L., Eikeset, A. M., McCauley, D. J., Payne, J. L., & Sunday, J. M. (2019). Greater vulnerability to warming of marine versus terrestrial ectotherms. Nature, 569(7754), 108–111.

R Core Team (2026). R: A Language and Environment for Statistical Computing. R Foundation for Statistical Computing, Vienna, Austria. URL https://www.R-project.org.

Reed, T. E., R. S. Waples, D. E. Schindler, J. J. Hard, and M. T. Kinnison. 2010. Phenotypic plasticity and population viability: the importance of environmental predictability. Proceedings of the Royal Society B: Biological Sciences 277:3391–3400.

Rohr, J. R., D. J. Civitello, J. M. Cohen, E. A. Roznik, B. Sinervo, and A. I. Dell. 2018. The complex drivers of thermal acclimation and breadth in ectotherms. Ecology Letters 21:1425–1439.

Ruel, J. J., and M. P. Ayres. 1999. Jensen’s inequality predicts effects of environmental variation. Trends in Ecology & Evolution 14:361–366.

Ruthsatz, K., F. Dahlke, K. Alter, S. Wohlrab, P. C. Eterovick, M. L. Lyra, S. Gippner et al. 2024. Acclimation capacity to global warming of amphibians and freshwater fishes: Drivers, patterns, and data limitations. Global Change Biology 30:e17318.

Scheiner, S. M. 1993. Genetics and evolution of phenotypic plasticity. Annual review of ecology and systematics 24:35–68.

Seebacher, F., C. R. White, and C. E. Franklin. 2015. Physiological plasticity increases resilience of ectothermic animals to climate change. Nature Climate Change 5:61–66.

Shah, A. A., B. A. Gill, A. C. Encalada, A. S. Flecker, W. C. Funk, J. M. Guayasamin, B. C. Kondratieff et al. 2017. Climate variability predicts thermal limits of aquatic insects across elevation and latitude. Functional Ecology 31:2118–2127.

Sheldon, K. S., and J. J. Tewksbury. 2014. The impact of seasonality in temperature on thermal tolerance and elevational range size. Ecology 95:2134–2143.

Sinclair, B. J., K. E. Marshall, M. A. Sewell, D. L. Levesque, C. S. Willett, S. Slotsbo, Y. Dong et al. 2016. Can we predict ectotherm responses to climate change using thermal performance curves and body temperatures? Ecology Letters 19:1372–1385.

Sunday, J. M., A. E. Bates, and N. K. Dulvy. 2011. Global analysis of thermal tolerance and latitude in ectotherms. Proceedings of the Royal Society B: Biological Sciences 278:1823–1830.

Vasseur, D. A., J. P. DeLong, B. Gilbert, H. S. Greig, C. D. Harley, K. S. McCann, V. Savage et al. 2014. Increased temperature variation poses a greater risk to species than climate warming. Proceedings of the Royal Society of London B: Biological Sciences 281:20132612.

von May, R., A. Catenazzi, A. Corl, R. Santa-Cruz, A. C. Carnaval, and C. Moritz. 2017. Divergence of thermal physiological traits in terrestrial breeding frogs along a tropical elevational gradient. Ecology and Evolution 7:3257–3267.

Weaving, H., J. S. Terblanche, P. Pottier, and S. English. 2022. Meta-analysis reveals weak but pervasive plasticity in insect thermal limits. Nature Communications 13:5292.

Wickham, H., Averick, M., Bryan, J., Chang, W., McGowan, L. D. A., François, R., … & Yutani, H. (2019). Welcome to the Tidyverse. Journal of open source software, 4(43), 1686.

Yilmaz, A. R., C. R. da Silva, and S. E. Diamond. 2026. Climatic variability drives greater thermal tolerance breadth and enables larger range size in butterflies. Biological Journal of the Linnean Society 147:blag003.

